# POSTERIOR-SUPERIOR INSULA REPETITIVE TRANSCRANIAL MAGNETIC STIMULATION REDUCES EXPERIMENTAL TONIC PAIN AND PAIN-RELATED CORTICAL INHIBITION IN HUMANS

**DOI:** 10.1101/2024.05.14.594260

**Authors:** Nahian S Chowdhury, Samantha K Millard, Enrico de Martino, Dennis Boye Larsen, David A Seminowicz, Siobhan M Schabrun, Daniel Ciampi de Andrade, Thomas Graven-Nielsen

## Abstract

High frequency repetitive transcranial magnetic stimulation (rTMS) to the posterosuperior insula (PSI) may produce analgesic effects. However, the neuroplastic changes behind PSI-rTMS analgesia remain poorly understood. The present study aimed to determine whether tonic capsaicin-induced pain and cortical inhibition (indexed using TMS-electroencephalography) are modulated by PSI-rTMS. Twenty healthy volunteers (10 females) attended two sessions randomized to active or sham rTMS. Experimental pain was induced by capsaicin administered to the forearm for 90 minutes, with pain ratings collected every 5 minutes. Left PSI-rTMS was delivered (10Hz, 100 pulses per train, 15 trains) ∼50 minutes post-capsaicin administration. TMS-evoked potentials (TEPs) and thermal sensitivity were assessed at baseline, during capsaicin pain prior to rTMS and after rTMS. Bayesian evidence of reduced pain scores and increased heat pain thresholds were found following active rTMS, with no changes occurring after sham rTMS. Pain (prior to active rTMS) led to an increase in the frontal negative peak ∼45 ms (N45) TEP relative to baseline. Following active rTMS, there was a decrease in the N45 peak back to baseline levels. In contrast, following sham rTMS, the N45 peak was increased relative to baseline. We also found that the reduction in pain NRS scores following active vs. sham rTMS was partially mediated by decreases in the N45 peak. These findings provide evidence of the analgesic effects of PSI-rTMS and suggest that the TEP N45 peak is a potential marker and mediator of both pain and analgesia.

## INTRODUCTION

High frequency repetitive transcranial magnetic stimulation (rTMS) involves the delivery of magnetic pulses over the brain and has been shown to be a promising, safe, and non-invasive treatment for pain [38; 68; 71]. A common target in rTMS treatments for pain is the primary motor cortex (M1) [38]. However, M1 rTMS shows a 25-50% reduction in pain intensity in only ∼50% of chronic pain patients [5; 46; 47]. Moreover, the effects of M1 rTMS on pain are believed to be mediated by functional alterations in extra motor areas [2; 40; 42]. As such, one strategy to improve the effects of rTMS on pain has been to explore non-M1 rTMS targets, with one of these being the posterior insular cortex (PSI) [17; 21; 27; 30; 37; 49], which plays a critical role in pain processing [39; 53]. In 10 healthy individuals, the effect of a single session of cTBS of the operculo-insular cortex demonstrated a decrease in heat pain sensitivity [49]. Likewise, another study reported decreased heat pain sensitivity and reduced clinical pain intensities after 5 repeated sessions of 10 Hz rTMS to PSI in 31 patients with chronic neuropathic pain [30]. This suggests PSI-rTMS can reduce pain severity and decrease heat pain sensitivity. While this is encouraging, the neuroplastic changes that occur during PSI-rTMS analgesia remain unknown.

One method of assessing pain and analgesia-related neuroplasticity has been to use single pulse TMS to measure changes in corticomotor excitability (CME) in response to painful stimuli and following rTMS [11; 12; 15; 24; 51]. However, this methodology limits the investigation of neuroplasticity to the motor system and is an indirect measure of cortical plasticity given it is confounded by activity of spinal and subcortical processes [13]. A novel method combining TMS with electroencephalography (EEG) allows for neuroplastic changes to be directly measured from multiple cortical regions [33] with excellent temporal resolution. When TMS is applied to M1 during concurrent EEG, several commonly observed peaks are detected in the TMS-evoked potential (TEP), with larger peaks occurring at 45 ms (N45) and 100 ms (N100) linked to stronger inhibitory (GABAergic) neurotransmission [9; 19; 20; 60; 61].

Recent studies using TMS-EEG suggest cortical inhibitory processes may be implicated in rTMS-induced analgesia [13; 23]. One study showed the amplitude of the frontocentral N45 and N100 TEP peak was increased in response to acute heat pain, with a larger increase in the frontocentral N45 associated with higher pain ratings [13]. Furthermore, 10Hz rTMS over dorsolateral prefrontal cortex (dlPFC) led to a decrease in TEP indices of GABAergic activity (N45 and N100) in people with major depression [74], while another study in healthy individuals showed a decrease in the amplitude of the frontocentral TEP negative peak indexing GABA (N100) following 10Hz dlPFC rTMS, with this decrease associated with an increase in pain thresholds [77]. Thus, it is possible that changes in pain perception following PSI-rTMS are mediated by changes in TEP peaks that index GABAergic activity.

The present study aimed to determine whether tonic experimental pain and cortical inhibition are modulated by PSI-rTMS. The study was conducted on healthy human participants, with active or sham PSI-rTMS applied during tonic capsaicin-induced pain, and TEPs and thermal pain sensitivity measures assessed before pain, during pain before rTMS, and during pain after rTMS. It was hypothesized that active PSI-rTMS would induce i) a reduction in capsaicin-induced pain intensity, ii) a decrease in pain sensitivity, and iii) a reduction in TEPs that index GABAergic activity (N45 and N100).

## METHODS

### Participants

This study was conducted at the Center for Neuroplasticity and Pain (CNAP), Aalborg University, Aalborg, Denmark. All procedures adhered to the Declaration of Helsinki, with written, informed consent obtained prior to study commencement. The study was approved by the local ethics committee (Videnskabsetiske Komite for Region Nordjylland: N-20210047). A sample size calculation was conducted (G*Power 3.1.9.7) based on available means/SDs reported in a previous study exploring the effects of 10 Hz rTMS on frontal TEPs that index GABA [74] and another study exploring the effects of PSI-rTMS on pain thresholds/pain intensity in healthy individuals [49]. We also used our previous work which reported correlation values between repeated measurements of TEP peaks before and after pain [13]. For the effect of rTMS on pain intensity (α = 0.05, β = 0.8, d = 1.82), a sample size of at least 5 individuals was required, while for TEPs (α = 0.05, β = 0.8, *r* = .9, d = 0.92 for N45 and d = 0.97 for N100) a sample size of at least 12 individuals was required. We opted for a higher sample size of 20 participants to improve statistical power.

Twenty healthy participants (10 females; aged 26.5± 4.6 years (mean ± SD) were recruited through online advertisement. Participants were excluded if they presented with any acute pain, had a history or presence of chronic pain, neurological, musculoskeletal, psychiatric or other major medical condition, were pregnant and/or lactating, or were contraindicated for TMS (e.g., metal implants in the head) as assessed using the Transcranial Magnetic Stimulation Adult Safety Screen questionnaire [64]. To further characterize the mental health profile and degree of catastrophic thinking related to pain, participants completed the following questionnaires: Beck-Depression Inventory [8], State-Trait Anxiety Inventory Pain Catastrophizing Scale [70] and Positive and Negative Affective Schedule [76].

### Experimental Protocol

This study used a cross-over, randomised, sham-controlled design. Participants attended two sessions (spaced ∼2-3 weeks apart) of either active or sham PSI-rTMS in a randomized sequence. Each session involved the administration of tonic pain induced by capsaicin applied to the right volar forearm for 90 minutes (Fig. 1). Left PSI-rTMS was delivered ∼50 minutes after capsaicin administration for 7.5 minutes. TEPs using combined TMS-EEG to M1 followed by thermal pain sensitivity were assessed three times: at baseline (pre-pain), during capsaicin pain prior to rTMS (pain pre-rTMS: between ∼30-45 minutes after capsaicin administration), and after rTMS (pain post-rTMS: between ∼65-80 minutes after capsaicin administration).

**Figure 1.**
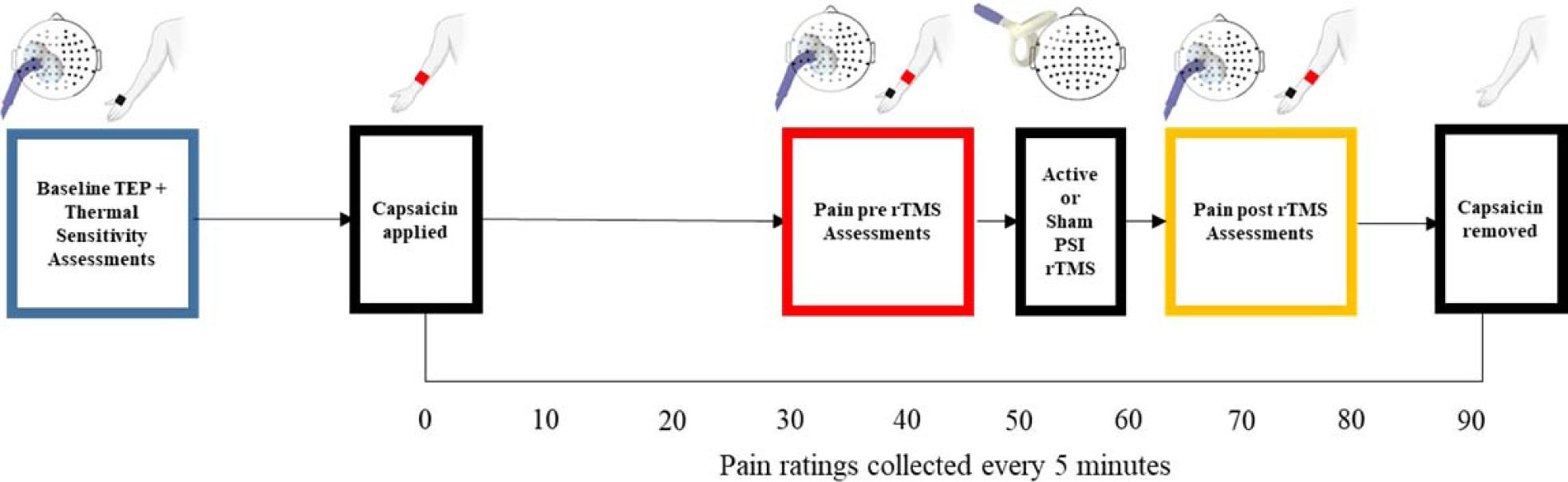
Diagram of the experimental protocol.

### Transcranial Magnetic Stimulation Evoked Electroencephalography

Participants sat in a comfortable chair with their eyes fixated on a cross placed on the wall in front of them. Single, biphasic transcranial magnetic stimuli were delivered using a Magstim unit (Magstim Ltd., UK) and 70 mm figure-of-eight flat coil. EEG was recorded using a TMS-compatible amplifier (g.HIamp EEG amplifier, g.tec-medical engineering GmbH, Schiedlberg, Austria) at a sampling rate of 4800 Hz. Signals were recorded from 63 passive electrodes, embedded in an elastic cap (EASYCAP GmbH, Etterschlag, Germany), in line with the 10-5 system. Recordings were referenced online to ‘right mastoid’ and the ground electrode placed on right cheekbone. This was to reduce artifacts produced by the TMS, which was applied on the left side of the head. Electrolyte gel was used to reduce electrode impedances below ∼5 kΩ. To maintain low impedances throughout the experiment, we used two net caps (GVB-geliMED GmbH, Ginsterweg Bad Segeberg, Germany) and a plastic stretch wrap handle film over the EEG cap [23]. In order to minimize the effect of the auditory response generated by the TMS coil click sound, a masking toolbox [66] was used with the participants wearing noise-cancelling headphones (Shure SE215-CL-E Sound Isolating, Shure Incorporated, United States).

Neuronavigation (Brainsight TMS Neuronavigation, Rogue Research Inc., Montréal, Canada) was used with a template MRI (MNI ICBM 152 average brain) from Brainsight software to track and calibrate each participant’s head position and TMS coil position in 3D space. Surface disposable silver/silver chloride adhesive electrodes (Ambu Neuroline 720, Ballerup, Denmark) were applied over the right first dorsal interosseous (FDI) muscle parallel to muscle fibres, with the ground electrode placed on the right ulnar styloid process. The coil was oriented at 45° to the midline, inducing a current in the posterior-anterior direction. To identify the left M1 target, the scalp site (‘hotspot’) that evoked the largest motor evoked potential (MEP) measured at the FDI was determined and marked. The rest motor threshold (RMT) was determined using the ML-PEST (maximum likelihood strategy using parametric estimation by sequential testing) algorithm to estimate the TMS intensity required to induce an MEP of 50 microvolts with a 50% probability [3]. This method has been shown to achieve the accuracy of methods such as the Rossini-Rothwell method [65] but with fewer pulses [69]. The test stimulus intensity was set at 90% RMT to minimize contamination of EEG signal from re-afferent muscle activation [13].

The real-time TEP visualization tool was used [10] to confirm that artefacts (muscle, auditory) in the signal were minimal, and that, given the coil orientation and 90% RMT stimulus intensity, that the early peaks (<100ms) at the stimulation site were evident (P30-N100) [10; 23; 43]. The neuronavigation system and real-time TEP visualization tool were used throughout each session to monitor coil positioning and TEP data quality across measurements within and between sessions. For each TEP measurement (baseline, pain pre-rTMS, pain post-rTMS), ∼150 TMS pulses (∼7 minutes total) were delivered with a jitter of 2.6-3.4 s [23; 43].

### Thermal Pain Sensitivity Assessment

Cold and heat pain thresholds were assessed at each timepoint (immediately after TEP measurement), in line with a previous study [13]. A 27 mm diameter thermode (Medoc Pathway ATS device; Medoc Advanced Medical Systems Ltd) was applied over the right thenar eminence. With the baseline temperature set at a neutral skin temperature of 32°C, participants completed two threshold tests in the following order: to report when a decreasing temperature first became painful (cold pain threshold, CPT) and to report when an increasing temperature first became painful (heat pain threshold, HPT). A total of three trials were conducted for each test to obtain an average, with an interstimulus interval of six seconds. Participants provided their threshold for each trial by pressing a button (with their left hand) on a hand-held device connected to the Medoc Pathway. Temperatures were applied with a rise/decrease rate of 1°C/s and return rate of 2°C/s (initiated by the button click).

### Heat-Evoked Pain

At each of the three timepoints, an additional test for heat-evoked pain was conducted. The heat thermode (Medoc Pathway ATS device; Medoc Advanced Medical Systems Ltd) was applied at the capsaicin administration site, and the temperature was increased towards a target temperature for 5 s (increase rate of 1°C/s, return rate of 2°C/s). The target temperature was determined at the baseline timepoint prior to capsaicin administration, by measuring the HPT (across 3 trials) over the capsaicin administration site, and then adding 2 degrees above this pain threshold. Participants provided a pain NRS rating to each 5 s stimulus, with this process repeated 3 times.

### Capsaicin-induced Tonic Pain

After the baseline TEP, heat-evoked pain and thermal sensitivity assessment, an 8% topical capsaicin patch (Transdermal patch, ‘Qutenza’, Astellas, 4× 4 cm) was applied to induce cutaneous pain over the volar part of the right forearm (5 cm from the wrist) [1]. A numerical rating scale (NRS) scale between 0 (no pain) to 10 (worst imaginable pain) was used to assess pain intensity every 5 minutes following patch application.

### Repetitive Transcranial Magnetic Stimulation

Active or sham rTMS over the orthogonal projection of the PSI was delivered with a double cone coil (D110, Magstim Ltd., UK). The location and intensity of stimulation was determined in between capsaicin administration and the first TEP measurement (i.e., within the first 30 minutes after capsaicin administration). The stimulation intensity was determined by delivering single pulse TMS using the double cone coil over the left motor representation of the tibialis anterior (TA) muscle, which has a similar depth within the cortex as the PSI [21]. The TA hotspot and RMT were determined by visually inspecting responses in the leg to the TMS pulse, with the RMT determined using the ML-PEST procedure [3].

The fast PSI method was used to identify the PSI target without the need for MRI-guided neuronavigation [18]. A recommended rTMS protocol for inducing analgesic effects was used: 1500 pulses (10 Hz, 15 trains of 10 s each, inter-train interval of 20 s, 7.5 minutes total) [16], with the intensity of stimulation set to 80% of the TA RMT [30], and the coil oriented so that the main phase of the biphasic waveform induced a current in the posterior-anterior direction [48]. In both active and sham conditions, an D70 figure-of-eight TMS coil (Magstim Ltd., UK) was placed in contact with the double cone coil but faced orthogonally using an adjustable mechanical arm. During active rTMS, the double cone coil was activated, whereas during sham rTMS, the second coil was activated [30].

### Data Processing

Pre-processing of the TEPs was completed using EEGLAB [25] and TESA [63] in MATLAB (R2021b, The Math works, USA), and based on previously described methods [13; 14; 55; 56; 63]. First, the data was epoched 1000 ms before and after the TMS pulse, and baseline corrected between −1000 ms and −5 ms before the TMS pulse. Bad channels which showed large decay artefacts from the TMS pulse were removed. The period between −5 ms and 12 ms after the TMS pulse was removed and interpolated by fitting a cubic function. Noisy epochs were identified via the EEGLAB auto-trial rejection function [26] and then visually confirmed. The fastICA algorithm with auto-component rejection was used to remove eyeblink and muscle artefacts [63]. The source-estimation noise-discarding (SOUND) algorithm was applied [55; 56], which estimates and supresses noise at each channel based on the most likely cortical current distribution given the recording of other channels. This signal was then re-referenced (to average). A band-pass (1-100 Hz) and band-stop (48-52 Hz) Butterworth filter was then applied. Any previously removed bad channels were then interpolated.

The grand-averaged TEPs (across participants) for the baseline, pain pre-rTMS, and pain post-rTMS were obtained. In line with previous studies investigating the effects of pain on TEPs [13], and rTMS on TEPs during pain [77], the mean TEP was extracted from a frontocentral region of interest (F1, F2, F3, F4, Fz, FC1, FC2, FC3, FC4, FCz). Peaks of the TEP from this ROI (e.g. N15, P30, N45, P60, N100, P180) were identified for each participant using the TESA peak function [63], with predetermined windows of interest (N15: 12-20 ms, P30: 25-40 ms, N45: 40-60 ms, P60: 55-70 ms, N100: 70-110 ms, P180: 150-200 ms) chosen to account for variation between participants in the latency of the peaks.

### Statistical Analysis

For CPT and HPTs, evoked pain NRS scores and TEP peak amplitudes, we computed the change (Δ) scores by subtracting the pain pre-rTMS timepoint and the pain post-rTMS timepoint from the baseline value of each respective session. Data are presented as mean ± standard deviations unless otherwise stated. Where relevant, Cohen’s *d* was reported to quantify effect sizes. Where violations of normality occurred according to Shapiro-Wilk tests, log-transformations of the data were conducted.

Bayesian inference was used to analyse the data, which considers the strength of the evidence for the alternative vs. null hypothesis, using JASP software (Version 0.12.2.0, JASP Team, 2020). Bayes factors were expressed as BF_10_ values, where BF_10_’s of 1–3, 3–10, 10–30, 30-100 and >100 indicated ‘weak’, ‘moderate’, ‘strong’, ‘very strong’ and ‘extreme’ evidence for the alternative hypothesis, while BF_10_’s of 1/3–1, 1/10-1/3, 1/30-1/10 and 1/100-1/30 indicated ‘anecdotal’, ‘moderate’, ‘strong’ ,‘very strong’ and ‘extreme’ evidence in favour of the null hypothesis [73]. Given the novelty of the study (no prior studies on PSI rTMS on TEPs), default priors in JASP were used to provide a balance between informed and non-informed hypotheses.

We first ran Bayesian paired t-tests to determine evidence for a difference between active and sham sessions in pain thresholds, evoked pain NRS scores and TEP peak amplitudes at the baseline timepoint. For capsaicin pain NRS Ratings, a 2 (session: active vs. sham) x 10 (timepoint: 0-45 minutes) Bayesian repeated measures ANOVA was conducted to determine the evidence for a difference in pain ratings between active and sham rTMS sessions prior to stimulation. Then, a 2 (session: active vs. sham) x 9 (timepoint: 50-90 minutes) Bayesian repeated measures ANOVA was conducted to assess pain NRS ratings following rTMS. The main effect of stimulation determined evidence for a difference in pain ratings between active and sham rTMS, while the interaction effect determined evidence for whether this difference changed across time. Follow-up Bayesian paired t-tests were conducted to compare pain ratings between the end of the session (90 minutes) and onset of rTMS (50 minutes) for each group separately.

For ΔHPT, ΔCPT, Δ evoked pain NRS scores and ΔTEP peak amplitudes, A 2 (session: active vs. sham) x 2 (timepoint: pain pre-RTMS, pain post-rTMS) Bayesian repeated measures ANOVA was conducted. The interaction between session and timepoint determined evidence of modulation of the outcomes as a result of active vs. sham rTMS. Follow-up Bayesian paired t-tests were conducted to determine evidence for a change in Δ scores between pain pre and pain post-rTMS for active and sham sessions separately. Note that the difference in the baseline-normalized Δ scores between pain pre and pain post-rTMS was identical to the difference in the non-normalized raw scores. Thus, increases and decreases in the Δ scores were directly interpreted as increases and decreases in the raw outcomes. Finally, a follow-up Bayesian one sample t-test was also conducted at each timepoint and session separately to determine overall change in outcomes relative to baseline.

For any TEP peaks and pain outcomes that demonstrated at least moderate evidence of a change between pre and post active rTMS, we further explored the link between these peak changes and both pain and analgesia. We conducted a Bayesian correlation analysis to determine whether, across the whole sample, ΔTEP peak amplitudes were associated with Δ pain outcomes following capsaicin administration (pain pre-rTMS – baseline) or following rTMS (pain post-rTMS – pain pre-rTMS). We also determined whether Δ pain outcomes following rTMS were mediated by ΔTEP peak amplitudes. We used the bmlm package in R [75] to conduct a mediation analysis for repeated measures designs. This package uses a Bayesian framework to compute the mean and 95% credibility interval of plausible posterior parameter values for the total effect of the mediation model, direct effects between each variable, and the indirect effect (i.e. mediation effect). For each model, the outcome variable was Δ pain outcome following rTMS (pain post-rTMS– pain pre-rTMS), the predictor variable was PSI-rTMS session (active vs sham) and the mediating variable was ΔTEP peak amplitude (pain post-rTMS – pain pre-rTMS).

## RESULTS

All participants completed the active and sham sessions, with no missing data. The mean interval between sessions was 21.5 ± 11.4 days. Eight out of 20 participants were able to correctly identify the sequence of the active and sham sessions, suggesting blinding was successful. All participants tolerated the rTMS without side effects. The mean scores on the questionnaires were 5.5 ± 8.1 for the Pain Catastrophizing Scale, 9.0 ± 6.2 for the Beck-Depression Inventory II, 25.4 ± 10.9 for the State-Trait Anxiety State Scale, 27.6 ± 9.4 the State-Trait Anxiety Trait Scale and 21.3 ± 6.0 and 7.1 ± 5.1 for the Positive and Negative Affect scales respectively. These mean scores do not indicate clinical levels of pain catastrophizing, depressive or anxiety symptoms [7; 45; 70].

The FDI RMT was 60.5 ± 8.6% for the active session and 60.0 ± 7.9% for the sham session, with moderate evidence for no difference between sessions (BF_10_ = 0.23). Similar to TA RMT values from previous studies using the double cone coil [28; 67], the TA RMT in the present study was 42.7 ± 6.0% for the active session and 43.0 ± 6.6% for the sham session, with moderate evidence for no difference between sessions (BF_10_ = 0.24).

### Capsaicin Pain NRS Ratings

Prior to rTMS (0-45 min post-capsaicin), there was extreme Bayesian evidence for an increase in pain NRS ratings following capsaicin administration (main effect of timepoint: BF_10_ = 6.4 x 10^32^, Fig. 2A), anecdotal evidence that pain NRS ratings did not differ between the active and sham rTMS sessions (main effect of session: BF_10_ = 0.57), and moderate evidence that this difference did not change across timepoints (session x timepoint interaction: BF_10_ = 0.15). After rTMS (50-90 min post-capsaicin), there was strong evidence for lower pain NRS ratings in the active versus sham rTMS session (main effect of session: BF_10_ = 25.4), and extreme evidence of this difference becoming larger over time (sessions x timepoint interaction: BF_10_ = 2.1×10^8^). Follow-up t-tests showed that, when comparing pain NRS ratings at 90 with 50 minutes, there was strong evidence that pain NRS reduced in the active rTMS session (BF_10_ = 14.30, *d* = -.76), and anecdotal evidence that pain NRS increased in the sham rTMS session (BF_10_ = 1.70, *d* = 0.50).

**Figure 2.**
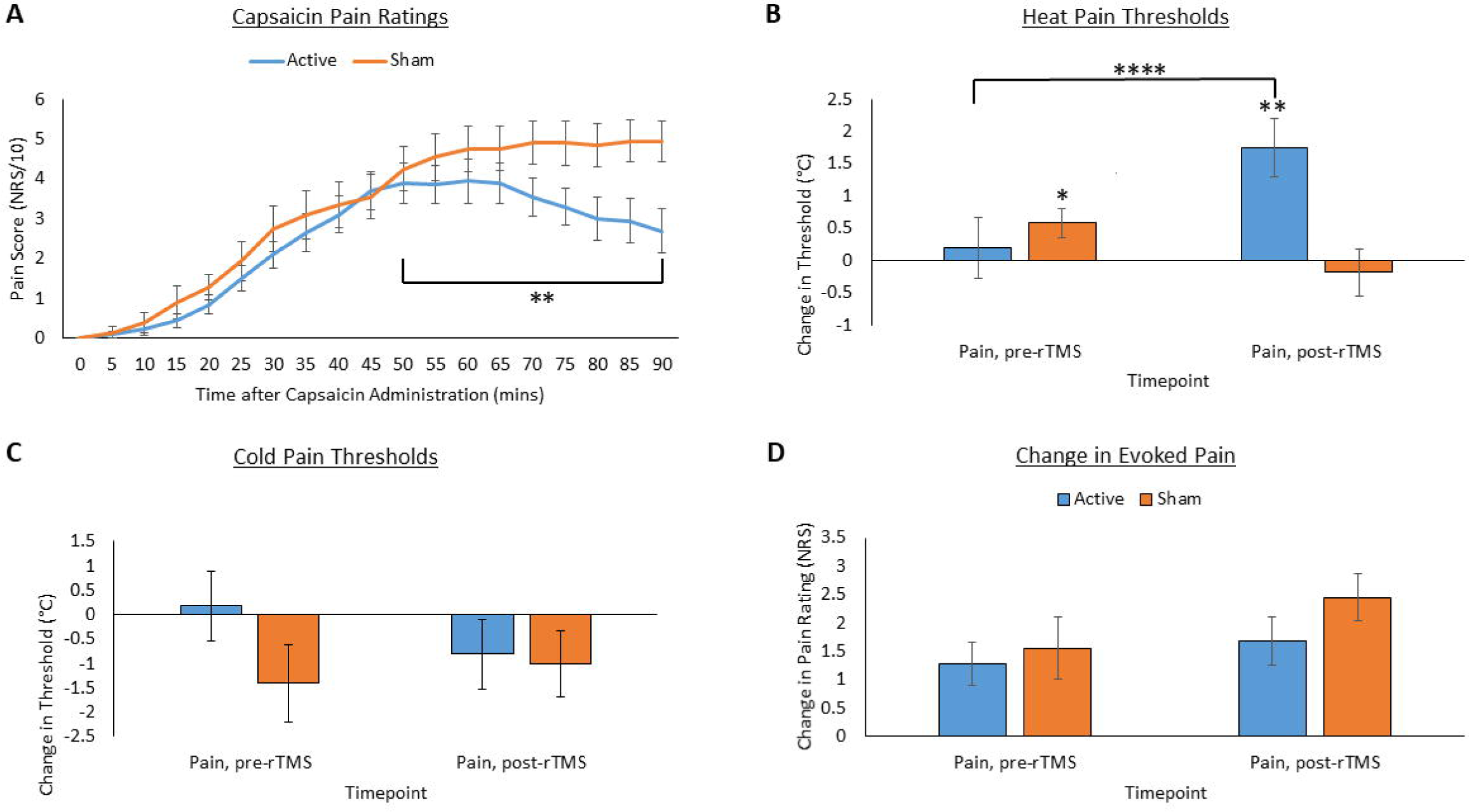
Mean (n = 20) and Standard Errors for capsaicin pain numerical rating scale (NRS) scores (**A**), Δ cold pain thresholds (**B**), Δ heat pain thresholds (**C**) and Δ evoked pain NRS scores (**D**) at each timepoint. *, **, ***, **** indicates moderate, strong, very strong and extreme Bayesian evidence of a difference between conditions or difference from baseline.

### Thermal Pain Sensitivity Distant to Capsaicin Application Site

Table 1 shows CPT, HPT and evoked pain scores at each timepoint before being normalized to baseline. There was anecdotal evidence for no difference in HPT (BF_10_ = 0.81) or CPT (BF_10_ = 0.45) between active and sham rTMS sessions at baseline. There was extreme Bayesian evidence that active rTMS modulated ΔHPT relative to sham (session x timepoint interaction: BF_10_ = 6419.72, Fig. 2B). Follow-up Bayesian paired t-tests showed that for the active rTMS session, there was extreme evidence that ΔHPT increased from the pain pre rTMS to pain post-rTMS timepoints (BF_10_ = 145080.16, *d* = 1.8). For the sham rTMS session, there was anecdotal evidence that ΔHPT decreased from pain pre-rTMS to pain post-rTMS timepoints (BF_10_ = 1.60, *d* = -.40). Follow-up Bayesian one sample t-tests showed that, at the pain pre-rTMS timepoint, there was moderate evidence that ΔHPT = 0 in the active session (BF_10_ = 0.25, *d* = .09) and moderate evidence that ΔHPT > 0 in the sham session (BF_10_ = 3.15, *d* = .58). At the pain post-rTMS timepoint, there was strong evidence that ΔHPT > 0 for the active session (BF_10_ = 40.28, *d* = 0.87), and moderate evidence that ΔHPT = 0 for the sham session (BF_10_ = 0.26, *d* = -.11). There was anecdotal evidence for no alteration in ΔCPT following active relative to sham rTMS (session x timepoint interaction: BF_10_ = 0.96, Fig. 2C).

**Table 1.** Thermal sensitivity outcomes at each timepoint for active and sham rTMS sessions before and after capsaicin-induced pain. CDT = Cold Detection Threshold, WDT = Warm Detection Threshold, CPT = Cold Pain Threshold, HPT = Heat Pain Threshold.

|  | Active rTMS |  |  | Sham rTMS |  |  |
| --- | --- | --- | --- | --- | --- | --- |
|  | Baseline | Pain<br>pre-rTMS | Pain<br>post-rTMS | Baseline | Pain<br>pre-rTMS | Pain<br>post-rTMS |
| **CPT (°C)** | 9.4±1.92 | 9.5±1.78 | 8.5 ±1.85 | 8.3±2.05 | 6.9±1.83 | 7.3±1.83 |
| **HPT (°C)** | 45.1±0.74 | 45.3±0.74 | 46.9 ±0.71 | 46.5±0.80 | 47.1 ±0.73 | 46.3 ±0.82 |
| **Evoked Pain**<br>**(NRS/10)** | 4.8±0.50 | 6.1±0.46 | 6.5±0.39 | 4.8±0.46 | 6.4±0.56 | 7.3±0.41 |

### Evoked Pain NRS Scores

Prior to capsaicin administration, the HPT at the target site was 44.1 ± 2.8 °C for the active session and 45.1 ± 3.3 °C for the sham session, with moderate evidence of no difference between sessions (BF_10_ = 0.24). There was moderate evidence of no difference in heat-evoked pain NRS scores between active and sham rTMS sessions at the baseline timepoint (BF_10_ = 0.23). There was anecdotal evidence that active rTMS did not modulate Δ evoked pain relative to sham (session x timepoint interaction: BF_10_ = 0.93, Fig. 2D).

### Transcranial Magnetic Stimulation Evoked Electroencephalography

Figures 3 and 4 show the grand-average TEPs and scalp topographies at each timepoint for the active and sham sessions respectively. Figure 5 shows the grand-average TEPs for the frontocentral ROI at each timepoint for the active and sham sessions. When comparing active and sham sessions at baseline, there was moderate evidence for no difference in the N15 (BF_10_ = 0.24), P30 (BF_10_ = 0.27), N45 (BF_10_ = 0.24), and P60 (BF_10_ = 0.31) peaks, anecdotal evidence for no difference in the P180 peak (BF_10_ = 0.45), and anecdotal evidence for a difference in the N100 peak (BF_10_ = 1.3).

**Figure 3.**
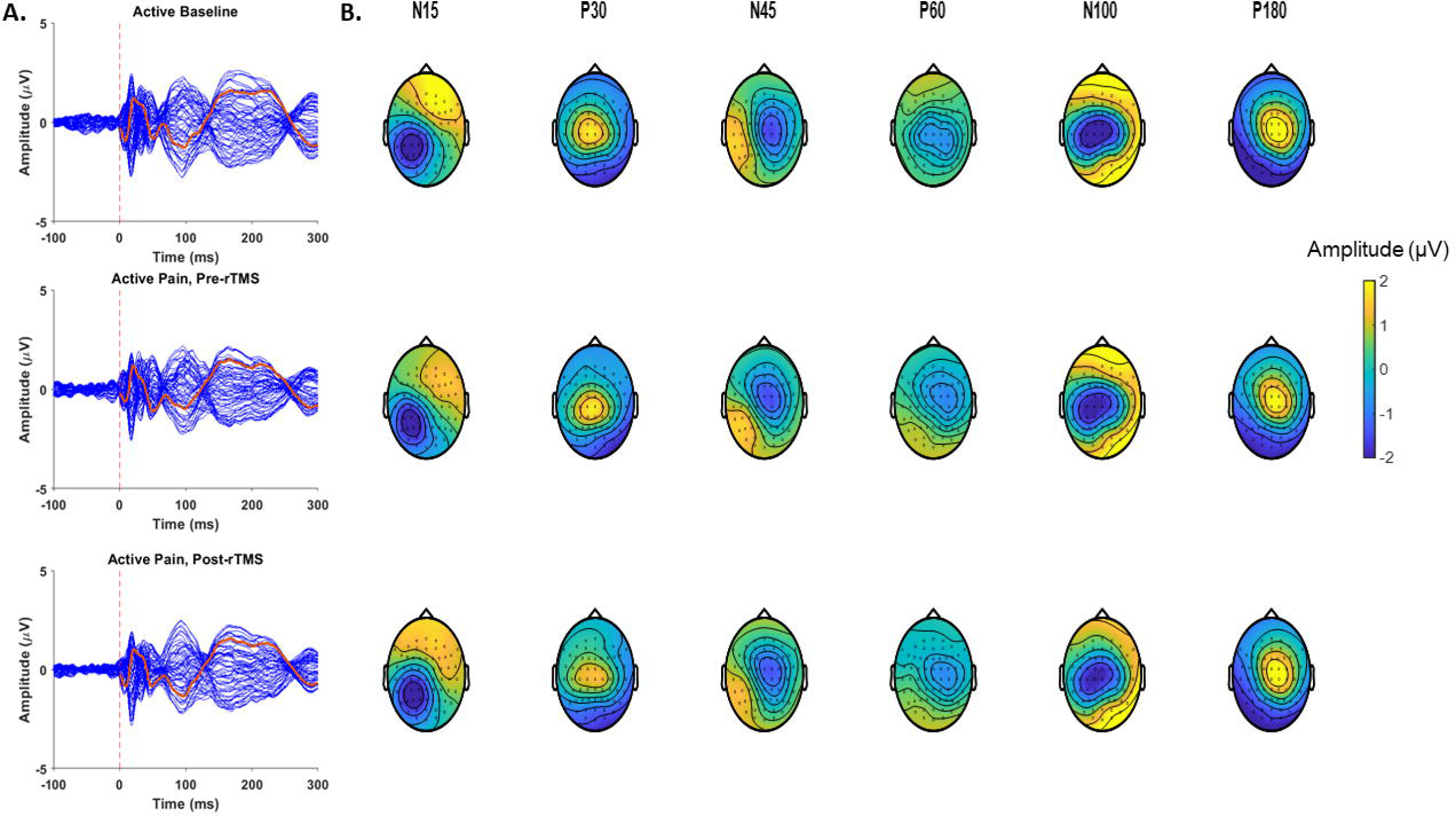
**A**: The Grand-average TEPs (n = 20) for all 63 channels during the baseline, pain pre-rTMS, and pain post-rTMS timepoints of the active repetitive transcranial magnetic stimulation session. The red line represents the mean TEP for the frontocentral region of interest. **B**: Scalp topographies and estimated source activity at timepoints where TEP peaks are commonly observed, including the N15, P30, N45, P60, N100, and P180.

**Figure 4.**
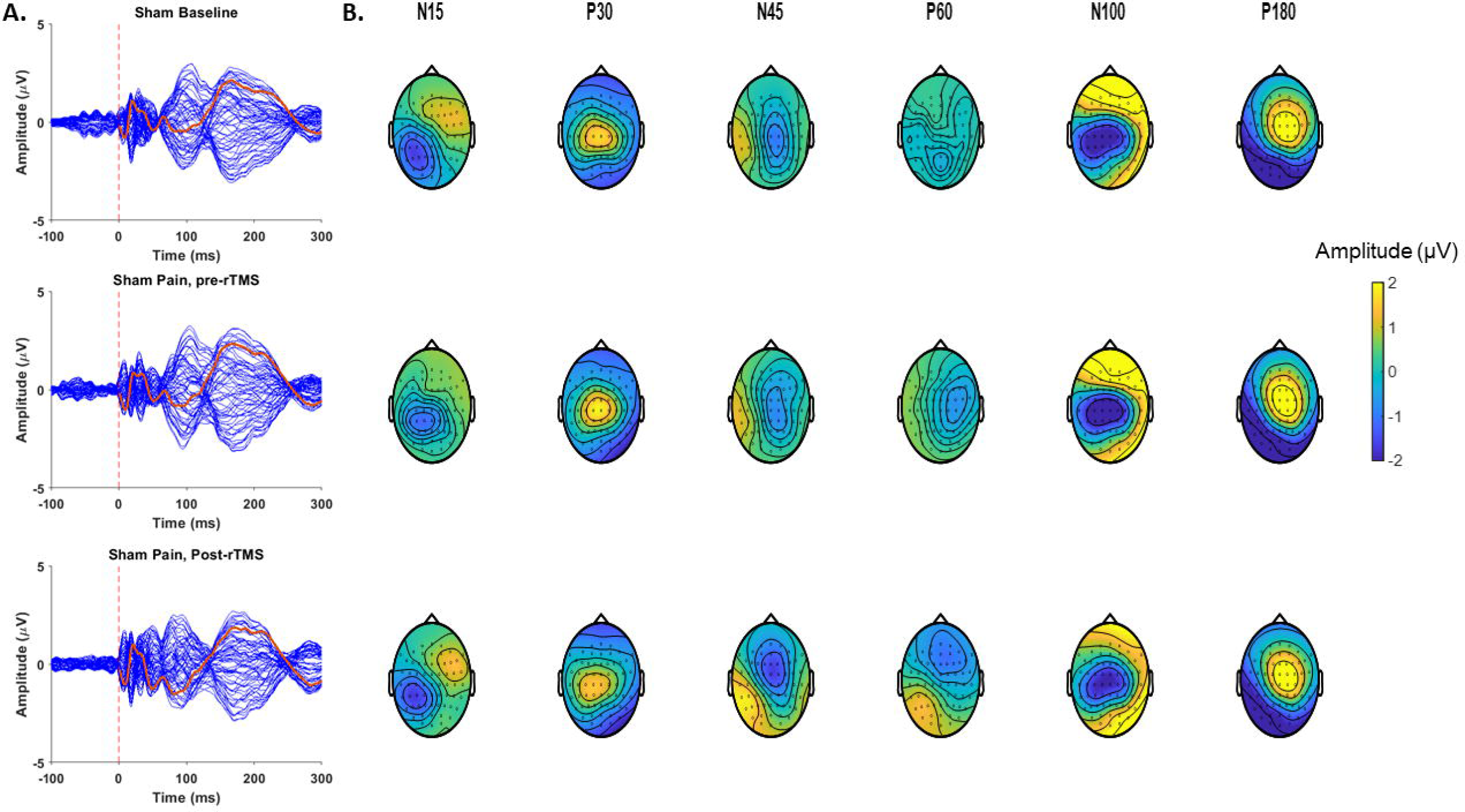
**A**: The Grand-average TEPs (n = 20) for all 63 channels during the baseline, pain pre-rTMS and pain post-rTMS timepoints of the sham repetitive transcranial magnetic stimulation session. The red line represents the mean TEP for the frontocentral region of interest. **B**: Scalp topographies and estimated source activity at timepoints where TEP peaks are commonly observed, including the N15, P30, N45, P60, N100, and P180.

**Figure 5.**
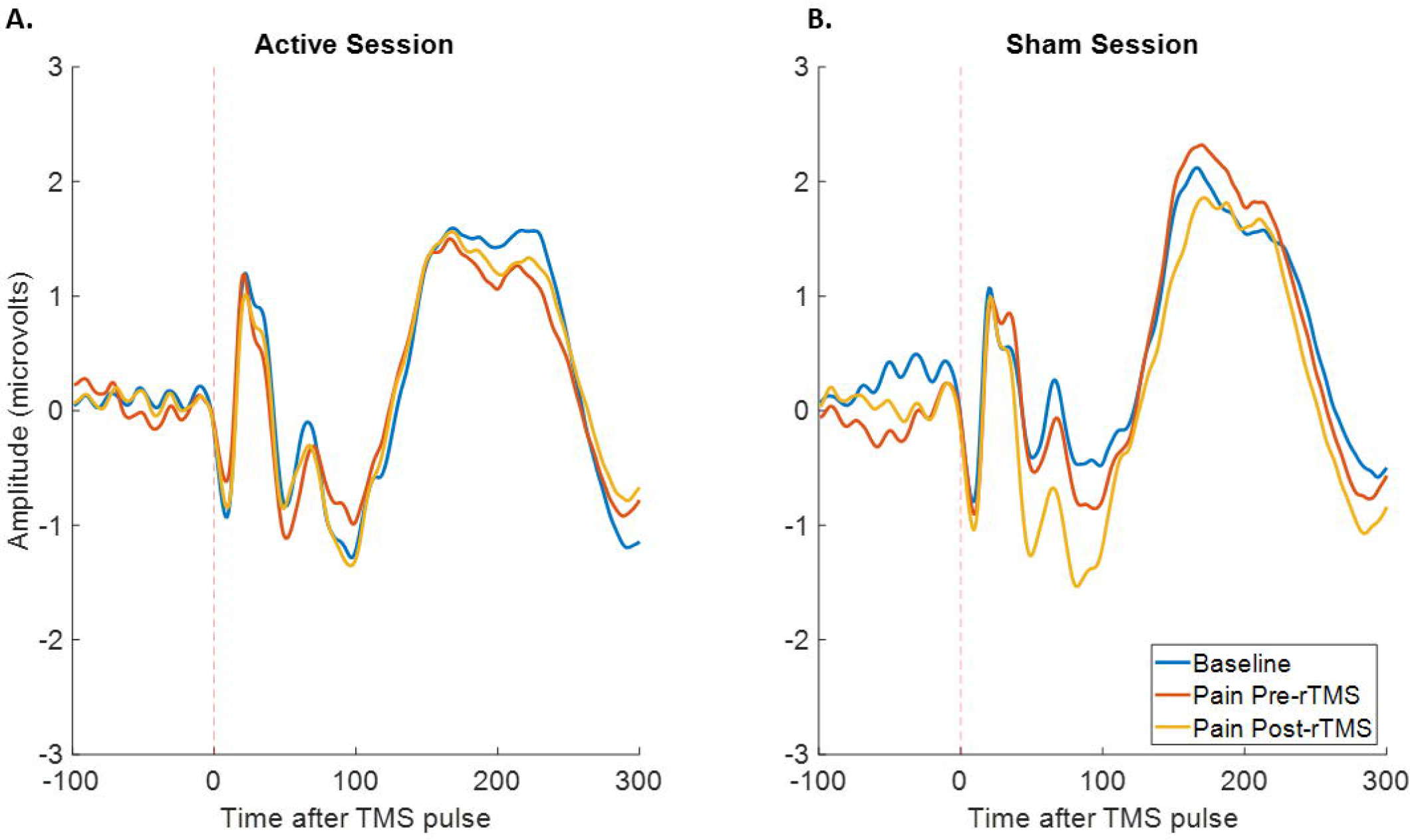
The Grand-average TEPs (n=20) for the frontocentral ROI during the baseline, pain pre-rTMS and pain post rTMS timepoints of the active **(A)** and sham (**B**) repetitive transcranial magnetic stimulation sessions.

Figure 6 shows the ΔTEP peak amplitudes (relative to baseline) at each timepoint for the active and sham sessions. There was strong evidence that active rTMS modulated ΔN45 relative to sham (session x timepoint interaction: BF_10_ = 18.65). Follow-up Bayesian t-tests showed moderate evidence that ΔN45 decreased from pain pre rTMS to pain post-rTMS timepoints following active rTMS (BF_10_ = 3.80, *d* = −0.60) and anecdotal evidence of no change in ΔN45 from the pain pre-rTMS to pain post-rTMS timepoints following sham rTMS (BF_10_ = 0.85, *d* = 0.39). A one sample t-test showed that at the pain pre-rTMS timepoint, there was moderate evidence ΔN45 < 0 in the active session (BF_10_ = 3.03, *d* = −0.57), and moderate evidence that the ΔN45 = 0 in the sham session (BF_10_ = 0.26, *d* = −0. 11). At the pain post rTMS timepoint, there was moderate evidence that ΔN45 = 0 for the active session (BF_10_ = 0.24, *d* = −0.07), and moderate evidence that ΔN45 < 0 for the sham session (BF_10_ = 3.02, *d* = 0.57). For all other peaks, there was no moderate evidence for any modulation by active rTMS relative to sham (BF_10_’s for all session x timepoint interactions < 3).

**Figure 6.**
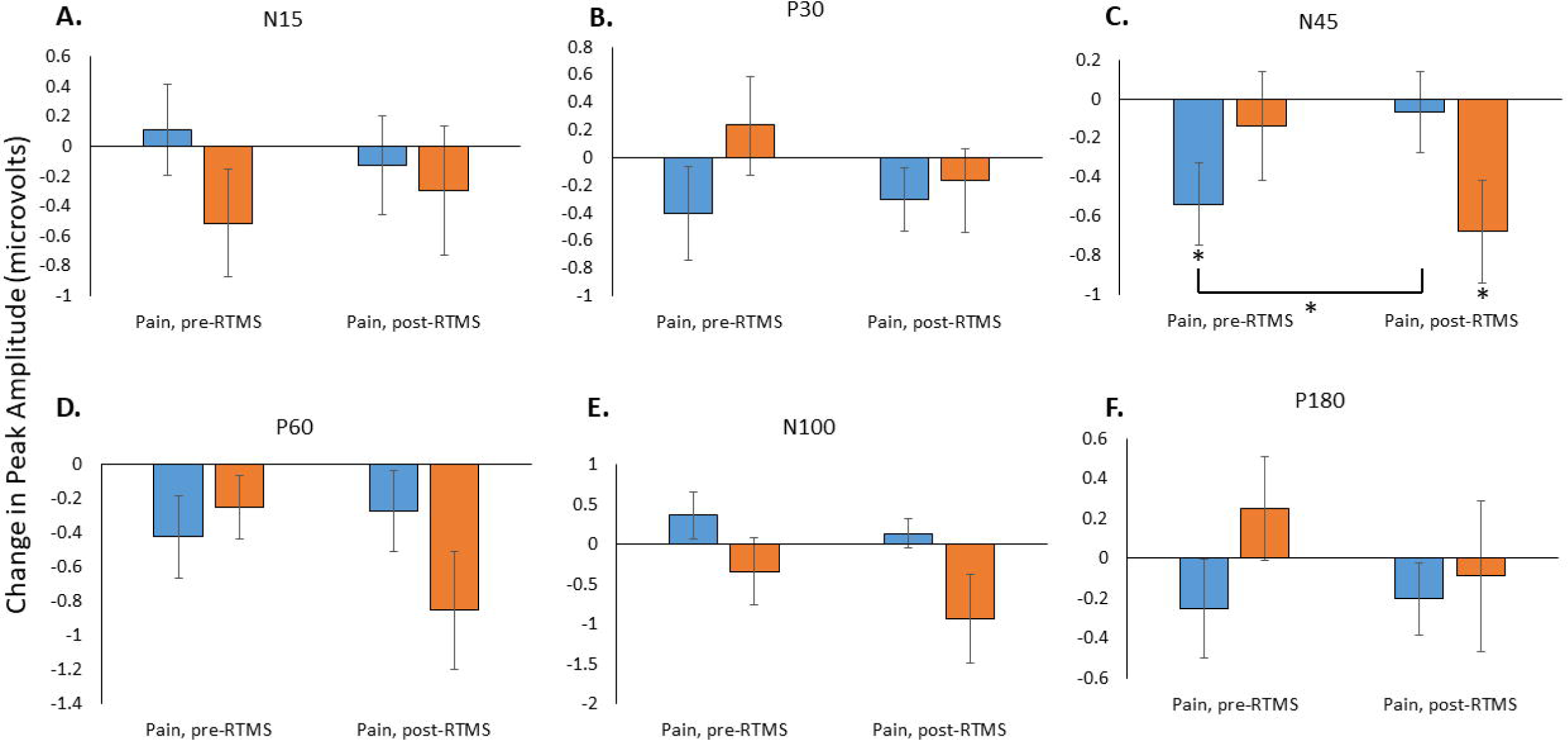
Mean (n = 20) and Standard Errors for ΔN15 (**A**), ΔP30 (**B**), ΔN45 (**C**), ΔP60 (**D**), ΔN100 (**E**) and ΔP180 (**F**) peaks normalized to baseline. * indicates moderate Bayesian evidence of a difference between conditions or a difference from baseline.

### Relationship between N45 peak changes and pain parameters following capsaicin administration

To further explore the link between ΔN45 and pain, we plotted the capsaicin pain NRS score at 45 minutes and ΔHPT at the pain pre-rTMS timepoint against ΔN45 at the pain pre-rTMS timepoint, pooled across both sessions (Figure 7A and 7B). There was moderate evidence for no correlation between ΔN45 and both capsaicin pain NRS score at 45 minutes (r_38_ = 0.03, BF_10_ = 0.20) and ΔHPT from baseline at the pain pre-rTMS timepoint (r_38_ = 0.11, BF_10_ = 0.24).

**Figure 7.**
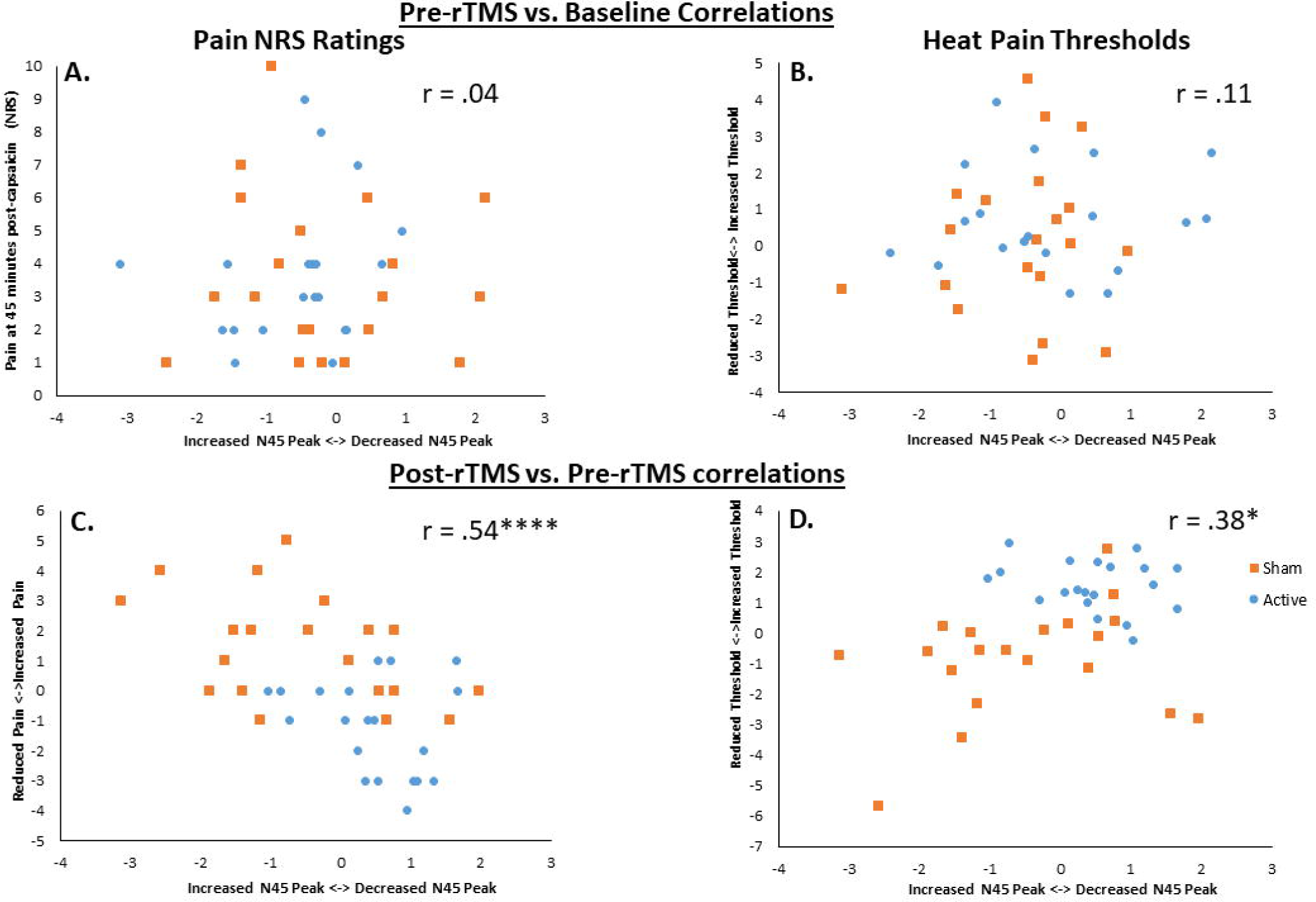
**(A)** Relationship between ΔN45 (pain pre-rTMS – baseline) and pain NRS at 45 minutes pooled across sessions (B). Relationship between ΔN45 (pain pre-rTMS – baseline) and Δ heat pain thresholds (pain pre-rTMS – baseline) pooled across sessions **(C)** Relationship between ΔN45 (pain post-rTMS – pain pre-rTMS) and Δ pain NRS (90– 50 mins). **(D)** Relationship between ΔN45 (pain post rTMS – pain pre rTMS) and Δ heat pain thresholds (pain post rTMS – pain pre rTMS). *, **, ***, **** indicates moderate, strong, very strong and extreme Bayesian evidence of a correlation.

### Relationship between N45 peak changes and pain parameters following rTMS

To explore the link between ΔN45 and reductions or increases in pain following rTMS, we plotted changes in pain NRS ratings (90 - 50 min) and changes in HPT (pain post-rTMS – pain pre-rTMS) against the change in the N45 peak (pain post rTMS – pain pre rTMS), pooled across sessions (Figure 7C and 7D). Across both sessions, we found extreme evidence that a larger reduction in capsaicin pain NRS scores were associated with a larger decrease in the N45 peak (r_38_ = 0.54, BF_10_ = 110.61), and moderate evidence that a larger increase in HPTs was associated with a larger decrease in the N45 peak (r_38_ = .38, BF_10_ = 3.4).

### Mediation Analysis

We determined whether the reductions in pain/increases in HPT following active vs. sham PSI-rTMS were mediated by decreases in the N45 peak. Two models were investigated (Fig. 8), where the outcome variable was Δpain-NRS (90 - 50 mins) or ΔHPT (pain post-rTMS – pain pre-rTMS), the predictor variable was PSI-rTMS session (active vs sham) and the mediating variable was ΔN45 (pain post-rTMS – pain pre-rTMS). When determining the total effect of PSI-rTMS on Δpain-NRS scores, the mean and credibility interval was 2.65 [1.59, 3.76] (i.e. a 2.65 stronger decrease in Δpain-NRS scores for the active vs. sham condition). The mean direct effect of rTMS on Δpain-NRS was 2.0 [0.9, 3.1] and the mean effect of rTMS on ΔN45 was −1.0 [0.9, 3.1] (i.e. a 1 µV stronger decrease in the ΔN45 in the active vs. sham condition). The mean indirect effect via ΔN45 was 0.65 [0.01, 1.61], suggesting 95% of plausible values of the indirect effect was above 0. This provides evidence that the reductions in pain by active vs. sham PSI-rTMS were partially mediated by decreases in the N45 peak. There was no evidence of mediation when analysing changes in HPT following rTMS.

**Figure 8.**
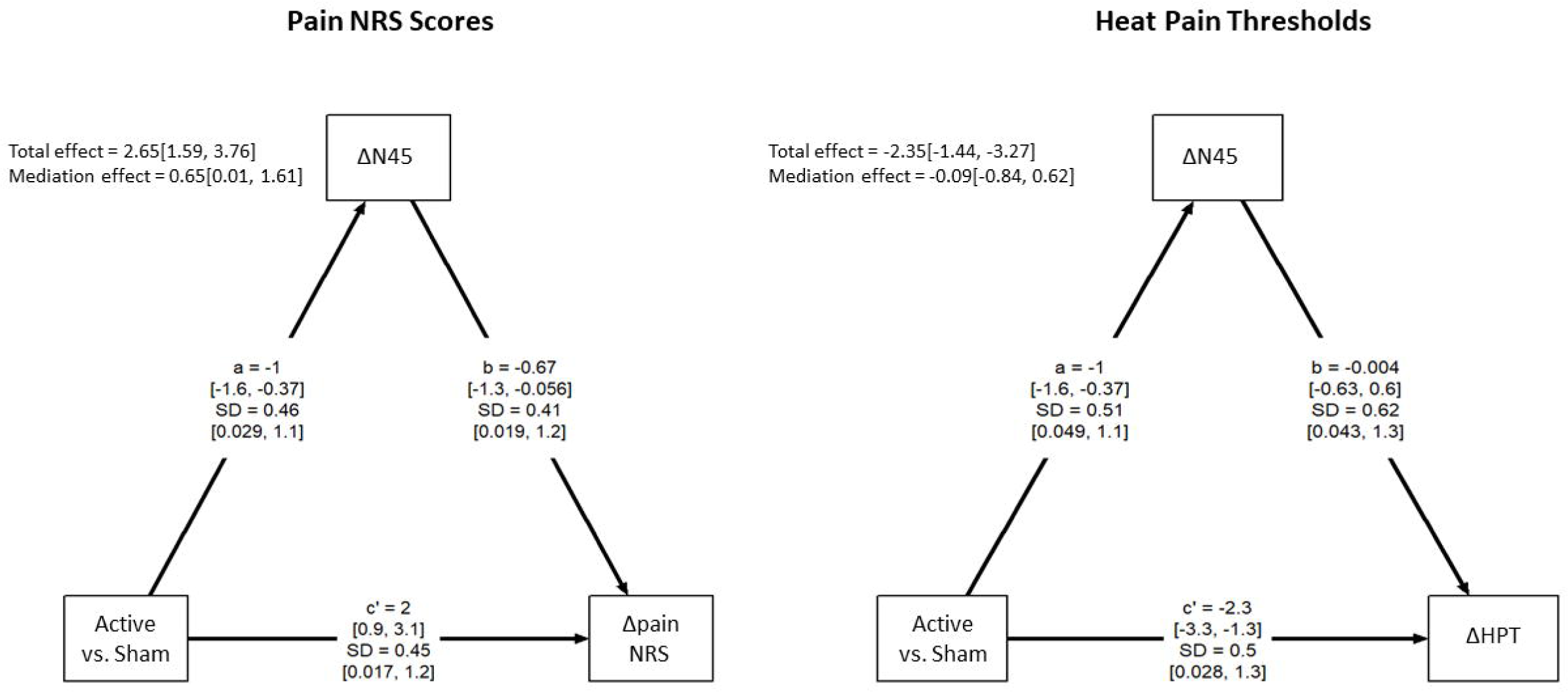
(A) Mediation model with session (active vs. sham) as the predicting variable, Δ pain NRS (90– 50 mins) as the outcome variable, and ΔN45 (pain post-rTMS - pain pre-rTMS) as the mediating variable. (B) Mediation model with session (active vs. sham) as the predicting variable, Δ heat pain thresholds (pain post-rTMS-pain pre-rTMS) as the outcome variable, and ΔN45 (pain post-rTMS -pain pre-rTMS) as the mediating variable. Credibility intervals for the mean effects and standard deviations are shown in brackets. In both models, the total effect represents the effect of session on the outcome in the absence of the mediator. The c’ path represents the direct effect of session on the outcome in the presence of the mediator, the a path represents the effect of session on ΔN45, and the b path represents the effect of ΔN45 on the outcome. The indirect effect determines the extent to which the effect of session on the outcome is accounted for by ΔN45.

## DISCUSSION

The present study aimed to determine whether tonic experimental pain and cortical inhibition assessed by TMS-EEG are modulated by PSI-rTMS. Active PSI-rTMS led to a decrease in capsaicin-induced pain intensity and increase in heat pain thresholds in body areas away from the site of experimental pain compared to sham. For the active rTMS session, pain prior to rTMS led to an increase in the frontal negative peak ∼45 ms (N45) TEP relative to baseline. Following active rTMS, there was a decrease in the N45 peak back to baseline levels. In contrast, the N45 was increased relative to baseline following sham PSI-rTMS. Lastly, decreases in pain intensity following active vs. sham rTMS were partially mediated by reductions in the N45 peak. Taken together, our findings suggest that 10Hz PSI-rTMS for ∼7.5 minutes has analgesic effects on experimental tonic pain. Furthermore, active PSI-rTMS not only leads to a decrease in cortical inhibition as indexed by the TEP N45 response, but these decreases partially mediate the effects of rTMS on pain intensity. Overall, these findings provide further insight into the role of cortical inhibitory processes during both pain and analgesia.

### Analgesic Effects of PSI-rTMS

The insular cortex is a key region involved in pain perception. The PSI receives sensory input from the spinal cord and thalamus [31], is activated during acute and chronic pain [6; 36], triggers painful sensations in response to direct electrical stimulation, and reduces painful sensations when lesioned [29]. The PSI is also believed to project descending inputs to GABAergic neurons within the brain stem. When PSI activity is increased, this triggers a loss of descending inhibition to the spinal cord leading to increased nociception [37]. rTMS is hypothesised to have a “blocking” effect on the PSI, which produces an antinociceptive/analgesic effect due to disinhibition of brain stem GABAergic neurons [37]. To determine whether PSI-rTMS does indeed have an antinociceptive/analgesic effect, we and others have determined the effects of PSI-rTMS using experimental pain models in healthy individuals and patients with neuropathic pain or epilepsy [17; 21; 27; 30; 37; 49]. The findings of the present study are largely consistent with previous studies, with a decrease in tonic pain intensity following active rTMS relative to sham and an increase in heat pain thresholds following active rTMS, although contrasted with no effects on cold pain thresholds or evoked heat pain NRS scores. In addition, this is the first study to demonstrate the effects of PSI-rTMS using a prolonged tonic experimental pain model (capsaicin), with previous experimental research using transient painful stimuli [49]. Furthermore, we demonstrate these analgesic effects for the first time using the fast PSI method [17], which precludes MRI-guided neuronavigation. New rTMS targets such as the PSI are actively being investigated as an alternative to M1 stimulation to increase the number of people with chronic pain responding to rTMS. However, targeting the PSI has traditionally required MRI guided neuronavigation, which can be time consuming and cost inefficient. The fast PSI method [18] was developed recently to reduce target identification time, and was shown to produce similar estimates of the PSI target compared to methods requiring neuronavigation, with high intra and inter-rater reliability. The finding that PSI-rTMS produced analgesia using the fast PSI method is promising for clinical application of the fast PSI method as this would greatly reduce time and costs required for targeted brain stimulation.

### Cortical Plasticity during PSI-rTMS analgesia

For the first time, we used TMS-EEG to investigate the potential neuroplastic changes during PSI-rTMS analgesia. In the active rTMS session, the TEP N45 peak was increased relative to baseline following capsaicin administration, which is consistent with previous work showing that tonic heat pain results in an increase in the N45 peak [13]. Following PSI-rTMS, the N45 peak was then decreased to baseline levels. This pattern of change in the N45 peak is consistent with previous work showing that larger increases in the N45 peak during tonic heat pain were associated with higher pain intensity [13]. In the sham session, we did not replicate an increase in the N45 peak at the pain pre rTMS point. The inconsistency might relate to the overall lower pain intensities at the pre-rTMS timepoint, with Figure 7A showing the majority of ratings were 3/10 or below which can be considered mild pain [72]. Indeed, in the sham rTMS condition, where pain gradually increased from the pre to post rTMS timepoint, the N45 peak was increased relative to baseline. Taken together, we argue that the natural response of the N45 peak is to increase in response to ongoing pain. When active rTMS is delivered, this tendency is reverted, with the N45 peak being brought back to baseline levels. These findings suggest that increased pain is associated with increases in the N45 peak, and analgesia is associated with decreases in the N45 peak.

To further unpack the association between the N45 peak and pain intensity, we determined whether individual changes in the N45 peak following active and sham rTMS were associated with increases or decreases in pain perception. As anticipated, across both sessions, increases in HPT and decreases in pain following rTMS were both associated with decreases in the N45 peak, suggesting that regardless of stimulation, the trajectory of pain correlates with the expected trajectory of the N45 peak. Further supporting the link between these measures, we found evidence that the reductions in pain NRS following active vs. sham rTMS were partially mediated by decreases in the N45 peak. This, for the first time, shows evidence for a potential causal role of the TEP N45 peak in the analgesic effects of rTMS. Currently, the mechanisms that mediate the analgesic effects of rTMS remain poorly understood [50]. While evidence suggests that PSI-rTMS leads to increased connectivity between cortical and subcortical structures directly involved in descending pain modulation [44; 52; 58], whether or not these alterations at the cortical or subcortical level, in turn, mediate the reductions in pain intensity is seldom investigated [50]. As such, our study provides crucial knowledge regarding these mediating mechanisms. This can inform targeted pain interventions, such that treatments that specifically reduce the TEP N45 peak may bring about larger pain reduction effect sizes.

Exactly how the analgesic effect of PSI-rTMS is mediated by the TEP N45 peak remains unclear. Source reconstruction showed that the TEP N45 peak might reflect activity within the sensorimotor cortex, despite its frontocentral topography in electrode space [13; 32]. Furthermore, pharmacological studies show that the TEP N45 peak reflects GABA_A_ receptor activity [60]. This suggests that the TEP N45 peak might reflect GABAergic activity within the sensorimotor cortex. Increased GABAergic activity in the sensorimotor cortex have been reported in response to painful thermal stimuli consistent with the present study [41]. Studies have shown that the primary and secondary somatosensory and motor cortices are functionally connected with the PSI, and that this connection is critical for the sensory discrimination aspect of pain processing [34; 35; 59]. Given PSI-rTMS is believed to block PSI function, this might result in downregulation of sensorimotor cortical GABA receptor activity resulting in antinociceptive effects. This hypothesis is speculative, as further multimodal work is required to elucidate the mediating role of TEP N45 peak on the analgesic effects of rTMS and the causal role of the TEP N45 peak in pain perception broadly. Further caution is also advised in interpreting the mediation analysis, given its exploratory nature and the relatively low sample size, and given a mediation of heat pain thresholds was not demonstrated.

The present study did not reveal alterations in other TEP peaks, such as the N100. This is inconsistent with research showing a decrease in the frontocentral N100 following 10 Hz rTMS to a different target, namely the dlPFC, with these decreases associated with increases in cold pain thresholds [77]. Beyond clear differences related to the targeting area, the effects on TEP peaks other than N45 may have occurred later, rather than immediately after TMS, given the analgesic effects of PSI-rTMS became larger overtime. Indeed, it has been suggested that the effects of M1 rTMS on pain seem to build up after at least the 1^st^ hour following stimulation [22; 57]. In this sense, it may have been suitable to record TEPs at multiple timepoints after rTMS to determine the onset and duration of effects.

### Strengths and limitations

This present study used a thorough experimental approach including a randomized sequence of sham and active sessions, separated by ∼2-3 weeks to minimize carry over effects, and TEP measurement based on real-time monitoring to improve data quality. Furthermore, successful blinding between active and sham sessions was achieved, and we reported no missing data. However, some limitations require attention. First, active rTMS over the PSI area can induce strong muscle activity in the temporalis and frontalis muscles. This may have, in turn, contributed to the analgesic effects as opposed to stimulation of the PSI per se. Future studies are encouraged to use control conditions that involve stimulation of the facial muscles. Another limitation is that visual inspection was used to estimate the motor threshold of the TA muscle instead of EMG. While this may have led to different estimation of the TA motor threshold, visual inspection has been shown to be a reliable method of RMT determination and some studies have shown no difference in RMT estimation between EMG and visual inspection [4; 62]. Moreover, the performance of visual methods is further improved when using ML-PEST [54], as was done in this study

### Conclusion

This study showed that PSI-rTMS reduces tonic experimental pain intensity and increases heat pain thresholds. These effects are accompanied by a decrease in cortical inhibition assessed by the TEP N45 response, with this decrease partially mediating the analgesic effects of rTMS. This study expands our understanding of the effects of PSI stimulation in humans showing that not only pain thresholds, but also experimental tonic pain is impacted by stimulating this target, and points to the N45 as a potential marker and mediator of analgesic effects of rTMS.

## Acknowledgements

Center for Neuroplasticity and Pain (CNAP) is supported by the Danish National Research Foundation (DNRF121). DCA is supported by a Novo Nordisk Grant (NNF21OC0072828). The present study was not pre-registered with an analysis plan

